# Nicotine e-cigarette vapor inhalation effects on nicotine & cotinine plasma levels and somatic withdrawal signs in adult male Wistar rats

**DOI:** 10.1101/833293

**Authors:** Christian Montanari, Leslie K Kelley, Tony M Kerr, Maury Cole, Nicholas W. Gilpin

**Affiliations:** Department of Physiology, School of Medicine, Louisiana State University Health Sciences Center, New Orleans, LA 70112, USA; La Jolla Alcohol Research Inc., La Jolla, CA, USA; Neuroscience Center of Excellence, Louisiana State University Health Sciences Center, New Orleans, LA 70112, USA; Alcohol & Drug Abuse Center of Excellence, Louisiana State University Health Sciences Center, New Orleans, LA 70112, USA; Southeast Louisiana VA Healthcare System (SLVHCS), New Orleans, LA, 70119, USA

**Keywords:** nicotine, ENDS, e-cigarette, vape, withdrawal, addiction

## Abstract

**Rationale:** Non-contingent chronic nicotine exposure procedures have evolved rapidly in recent years, culminating in Electronic Nicotine Delivery Systems (ENDS or e-cigarettes) to deliver vaporized drugs to rodents in standard housing chambers.

**Objectives:** The aim of the current work was to use ENDS to test concentration-dependent effects of nicotine e-cigarette vapor inhalation on blood-nicotine concentrations, blood-cotinine concentrations, and somatic withdrawal signs over time in rats.

**Methods:** Male Wistar rats were exposed to vapor containing various concentrations of nicotine (20, 40, 80 mg/mL) for 11 days through ENDS, and blood concentrations of nicotine and cotinine, the major proximate metabolite of nicotine, as well as spontaneous and precipitated somatic withdrawal signs were measured over time (across days of exposure and over hours after termination of vapor exposure).

**Results:** Exposing male Wistar rats to non-contingent nicotine vapor inhalation through ENDS produces somatic withdrawal symptoms and measurable blood-nicotine and blood-cotinine levels that change according to 1) concentration of nicotine in vape solution, 2) number of days of nicotine vapor exposure, 3) time since termination of nicotine vapor exposure, and 4) relative to the withdrawal signs, whether withdrawal was spontaneous or precipitated (by mecamylamine).

**Conclusions:** The data presented here provide parameters that can be used as a reasonable starting point for future work that employs ENDS to deliver non-contingent nicotine vapor in rats, although many parameters can and should be altered to match the specific goals of future work.

## Introduction

Electronic Nicotine Delivery Systems (ENDS or e-cigarettes) are battery-operated devices that people use to inhale nicotine. E-cigarettes are a very popular form of tobacco use among youth (Sing et al. 2015) including many who have never used conventional cigarettes (E-Cigarette Use Among Youth And Young Adults: A Report of the Surgeon General — Executive Summary.; 2016.), and are often advertised as a smoking cessation aid. However, adolescents with past e-cigarette use are more likely to smoke cigarettes later (Bold et al. 2017; Chaffee et al. 2017; Leventhal et al. 2017), and there is no conclusive evidence supporting the advertised effectiveness of ENDS for long-term smoking cessation. In fact, e-cigarettes facilitate cigarette smoking in adults (Kulic et al. 2018), and only nine percent of people that quit smoking cigarettes use ENDS as a smoking cessation aid (Weaver et al. 2018). Moreover, the safety of ENDS has not been extensively evaluated (Kitzen et al. 2019). This is particularly important considering that ENDS liquid solutions contain a variety of chemicals that are inhaled during the vaporization process (Sleiman et al. 2016; Hess et al. 2017).

Traditional animal models of nicotine dependence have used subcutaneous osmotic mini-pumps to deliver non-contingent chronic nicotine to rodents and intravenous (i.v.) catheterization to allow animals to voluntarily self-administer lower doses of nicotine (see Cohen et al. 2013 for a critical review). Osmotic mini-pumps do not mimic voluntary drug intake, i.v. self-administration produces weak withdrawal signs often without escalation of drug intake, and neither model mimics the route of administration seen in humans (see Cohen et al. 2013 for a critical review). Furthermore, the nicotine solution used in those types of studies mimics neither cigarettes nor e-cigarette vapor solution. To overcome these limitations, an aerosolization system was developed to pass pure air through nicotine liquid, which resulted in nicotine air that could be delivered chronically to rodents living in standard housing cages (George et al 2010; Cohen et al. 2013; Gilpin et al. 2014). This model produces blood-nicotine levels similar to those observed in human smokers (George et al. 2010; Matta et al. 2013; Gilpin et al. 2014), somatic signs of withdrawal upon termination of chronic nicotine exposure (George et al. 2010; Gilpin et al. 2014) and escalation of intravenous nicotine self-administration in nicotine vapor-exposed rats (Gilpin et al. 2014). However, this approach also does not use the same liquid solution or electronic and heat elements that are contained in e-cigarettes used by humans.

E-cigarettes consist of a battery-powered heating element, a cartridge with a liquid solution containing nicotine, flavorings, and other chemicals, and a mouthpiece to inhale. Recently, we began using ENDS to deliver vaporized drugs to rodents in standard housing chambers. This method has been used to administer vaporized Δ9-tetrahydrocannabinol (THC; Nguyen et al. 2016a, 2018; Javadi-Paydar et al. 2018, 2019), opiates (Vendruscolo et al. 2018), stimulants (Nguyen et al. 2016b, 2017), nicotine (Javadi-Paydar et al. 2019), or combinations of these drugs (Javadi-Paydar et al. 2019). ENDS nicotine inhalation produces behavioral and physiological effects in rats, such as increases in spontaneous locomotor activity and decreases in body temperature (Javadi-Paydar et al. 2019). Here, we chronically exposed male Wistar rats to different concentrations of nicotine, then we measured nicotine and cotinine (the major proximate metabolite of nicotine [Benowitz 1984] often used as a biomarker for nicotine exposure [Perez-Stable et al. 1995]) blood concentrations, and physical (spontaneous and precipitated) withdrawal symptoms over time (between and within sessions). Our aim was to use ENDS to test the concentration-dependent effects of nicotine on blood-nicotine concentrations, blood-cotinine concentrations, and somatic withdrawal signs over time in rats.

## Methods

### Animals

Adult male Wistar rats (N=18) (Charles River, Raleigh, NC, USA) weighing ~250 g at the time of arrival were pair-housed in a humidity- and temperature-controlled (22°C ±1) vivarium on an inverted 12 h light/dark cycle (lights off at 8 a.m.), with *ad libitum* access to food and water except during the experimental sessions. Rats were acclimated and handled for 1 week before the start of the experiments. Behavioral tests occurred during the dark phase. All procedures were approved by the Institutional Animal Care and Use Committee of the Louisiana State University Health Sciences Center and were in accordance with the National Institute of Health Guidelines.

### Drugs

(−)-Nicotine free base (Cat # N3876, Sigma-Aldrich, St. Louis, MO, USA) was mixed with a 50:50 propylene glycol (PG) and vegetable glycerin (VG) blend (Fisher Scientific, Pittsburgh, PA, USA) for aerosol administration (PG:VG ratio was selected based on the fact that e-cigarette liquid formulations used by humans are often composed of 50:50 PG:VG (Peace at al. 2016). Mecamylamine hydrochloride (Cat # M9020, Sigma-Aldrich), a non-selective and non-competitive antagonist of the nicotinic acetylcholine receptors (nAChRs) (Bacher et al. 2009) was dissolved in saline and administered via subcutaneous (s.c.) injections.

### Apparatus

Six sealed exposure chambers were modified from the Allentown, Inc (Allentown, NJ, USA) standard rat cages (259mm × 234mm × 209mm) to deliver vaporized nicotine into the chambers (see Figure 1). Six e-Vape generator controller boxes (Model SVS1; La Jolla Alcohol Research, Inc., La Jolla, CA, USA) set at 4.5 watts were triggered via MedPC IV software (Med Associates, St. Albans, VT, USA) to deliver to each chamber nicotine vapor puffs from e-cigarette cartridges (Protank II Atomizer, MT32 coil operating at 2.2 ohms; Kanger Tech; Shenzhen Kanger Technology Co., LTD; Fuyong Town, Shenzhen, China). Chamber air was vacuum controlled to set the air flow rate at ~3.5 L/min. Once generator controllers were triggered, nicotine vapor was integrated with the ambient air stream and delivered to the chambers via polyethelene tubing connected to the cartridges. Nicotine vapor was pulled from the chamber to an exhaust via vacuum air pumps connected to the chambers (see Figure 1).

**Fig. 1.**
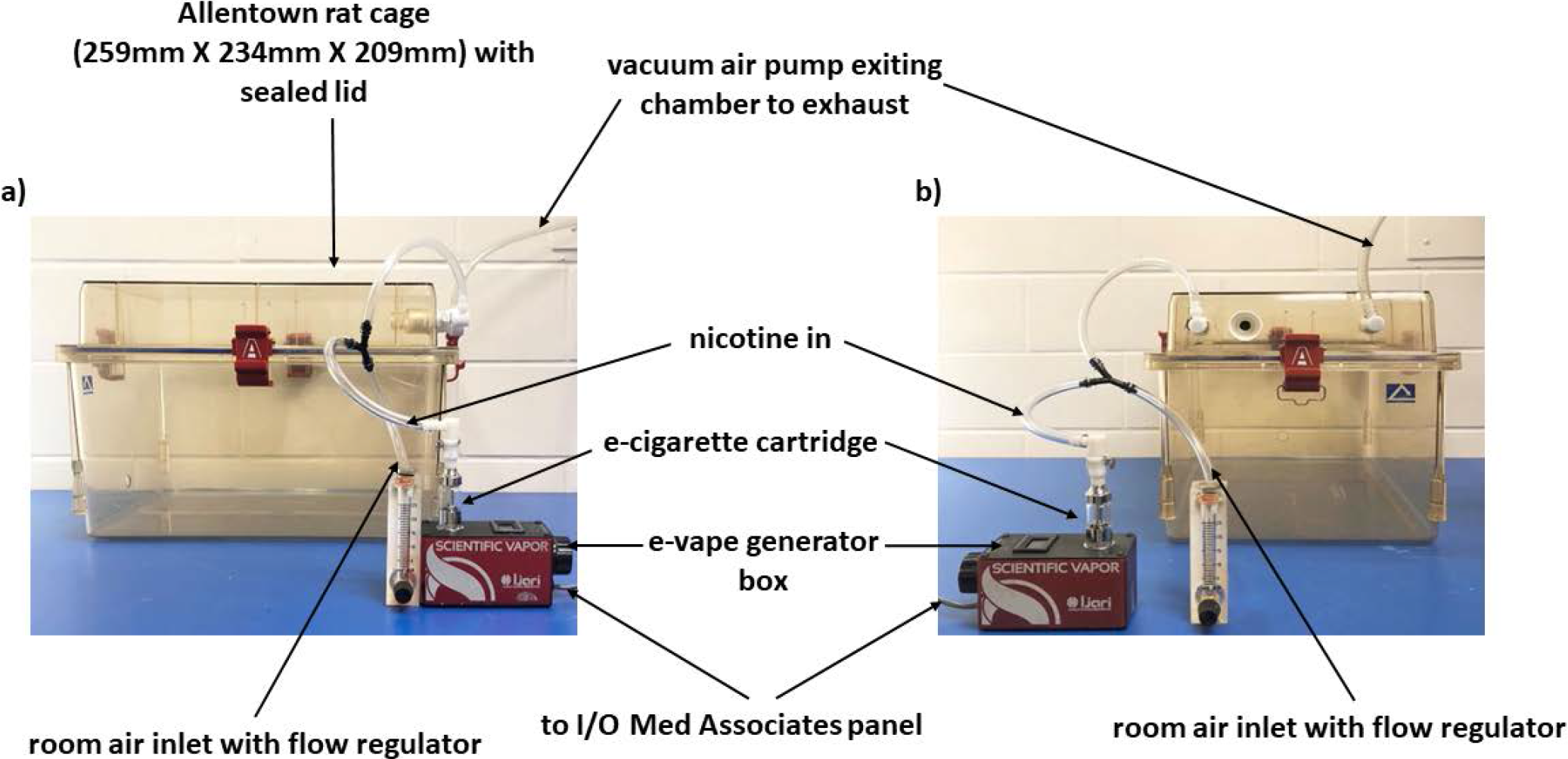
Front (**panel a**) and side (**panel b**) view of the apparatus used

### Experimental procedure

#### Nicotine Vapor Exposure

Rats were randomly divided into three groups (N=6/group) that were each exposed to one of three vaporized nicotine concentrations (20, 40 or 80 mg/mL) for 11 consecutive days (60 min/session/day; one 3-s nicotine puff delivered every 2 minutes for a total of 30 puffs over 60 minutes (all parameters chosen based on pilot studies conducted in our lab). The animals were run in three separate cohorts of 6 subjects (in each cohort there were 2 rats for each concentration dose). Chamber assignments were counterbalanced between cohorts and dose groups. All sessions were run 3-4 hours after the start of the dark phase. Concentrations for nicotine vapor administration are expressed as mg/mL in the e-cigarette cartridges. The e-cigarette cartridges here used can hold up to 2.5 ml; around 1 ml of nicotine solution was required in a 60-minute session. Nicotine concentrations were selected based on three factors: 1) the nicotine concentration (0-30 mg/mL) usually found in commercial e-cigarette liquids (Goniewicz et al. 2013); 2) JUUL and other companies have recently popularized nicotine salt solutions with nicotine levels in e-cigarette liquids up to 60-100 mg/ml; 3) rapid metabolism of nicotine in rats compared to humans (t_1/2_=45 min in rats vs 2 h in humans) necessitate higher nicotine doses in rat studies that aim to produce blood-nicotine concentrations similar to those seen in human smokers (see Matta et al. 2006 for a review).

#### Plasma nicotine and cotinine measurement

On days 1 and 10, blood samples were collected from each rat at multiple time points in series (0, 15, 30, 60 and 120 minutes after termination of nicotine exposure) to measure nicotine and cotinine plasma levels. Tail blood samples (~500 μL) were collected by making a cut 1 mm from the tip of the tail; the same cut was used for all subsequent bleeds of each animal. Blood from each animal was collected in Eppendorf tubes and kept on ice. After centrifugation, plasma (~50 μL) was stored at −80°C until analysis. Plasma nicotine and cotinine content was analyzed and quantified by an experimenter blinded to the experimental conditions, using liquid chromatography/mass spectrometry (LCMS). Briefly, 50 μl of plasma were mixed with 50 μl of deuterated internal standard (100 ng/ml cotinine-d3 and nicotine-d4; Cerilliant). Nicotine and cotinine (and the internal standards) were extracted into 900 μl of acetonitrile and then dried. Samples were reconstituted in 100 μL of an acetonitrile/water (9:1) mixture. Separation was performed on an Agilent LC1200 with an Agilent Poroshell 120 HILIC column (2.1mm × 100mm; 2.7 um) using an isocratic mobile phase composition of acetonitrile/water (90:10) with 0.2% formic acid at a flow rate of 325 μL/min. Nicotine and cotinine were quantified using an Agilent MSD6130 single quadrupole interfaced with electrospray ionization and selected ion monitoring [nicotine (m/z=163.1), nicotine-d4 (m/z=167.1), cotinine (m/z=177.1) and cotinine-d3 (m/z=180.1)]. Calibration curves were generated daily using a concentration range of 0-200 ng/mL with observed correlation coefficients of 0.999. Plasma nicotine and cotinine levels were separately analyzed using three-way ANOVAs with group (three levels of nicotine concentration: 20, 40 and 80 mg/mL) as a between-subjects factor, and day (two levels: day 1 *vs* day 10) and time (5 levels: 0, 15, 30, 60 and 120 minutes post-nicotine exposure) as within-subject factors.

#### Spontaneous and mecamylamine-induced withdrawal assessment

Rats were evaluated by a blind observer for both spontaneous and mecamylamine-precipitated withdrawal as previously described (Malin et al. 1992, 1994; George et al. 2010; Gilpin et al. 2014). Briefly, rats were placed into a clean cage and scored over 10 minutes for number of occurrences of the following somatic signs: eye blinks, head shakes, yawn, body shakes, ptosis, teeth chatter, tremors, scratch/bite, grooming/licking. Repeated successive counts of any sign required a distinct pause between episodes. For each rat, the total number of somatic signs was summed for an overall withdrawal (WD) score. Rats were initially scored the day before the first day of nicotine exposure (baseline condition). Spontaneous withdrawal signs were then evaluated 24 hours after the first day of nicotine exposure (day 1 at 24 hours post-nicotine), and again on day 10 at 1, 2, 4, 6 and 24 hours post-nicotine exposure. On day 11, one hour following termination of nicotine exposure, rats were injected with mecamylamine (1.5 mg/kg, s.c.) and precipitated withdrawal was assessed at 1, 2, 4, 6, 24 and 48 hours post-nicotine exposure (at 0, 1, 3, 5, 23 and 47 hours post-mecamylamine injection).

The baseline WD score was first compared with the WD score obtained 24 hours post-nicotine exposure at day 1 and 10 (spontaneous withdrawal) and 11 (precipitated withdrawal), via a 2-way ANOVA with group (three levels of nicotine concentration: 20, 40 and 80 mg/mL) as between-subject factor and session (4 levels: baseline, day 1, 10 and 11) as within-subject factor. Subsequently, baseline WD scores were compared with spontaneous withdrawal symptoms at day 10 using a 2-way ANOVA with group as between-subject factor and time (6 levels: baseline and 1, 2, 4, 6 and 24 hours post-nicotine exposure at day 10) as within subject factor. Precipitated withdrawal symptoms were also analyzed using a 2-way ANOVA with group as between-subject factor and time (7 levels: baseline and 1, 2, 4, 6, 24 and 48 hours post-nicotine exposure at day 11) as within-subjects factor. For each group and specific withdrawal time point, we also analyzed differences between spontaneous withdrawal symptoms (i.e., day 10) and precipitated withdrawal symptoms (i.e., day 11) using 2-tailed paired t-tests.

### Statistical analysis

Statistical analyses were performed using IBM SPSS Statistics 25 software. All variables are expressed as mean number ± SEM. Significant main or interaction analysis of variance (ANOVA) effects (p ≤ 0.05) were followed by Bonferroni post-hoc tests, which adjusted p-value significance criteria for multiple comparisons. Graphs were generated using GraphPad Prism 7.05 software.

## Results

### Plasma-nicotine levels

Analysis of plasma-nicotine levels yielded significant main effects of time [F(3.141,60)=56.710, p<0.001) and group [F(2,15)=74.743, p=0.001), a significant time × group interaction effect [F(4,60)=9.354, p=0.001), a significant day × time interaction effect [F (4,60)=4.447, p=0.003], and a significant day × time × group interaction effect [F(8,60)=4.618, p=0.001). As shown in Figure 2a, Bonferroni post-hoc tests confirmed that on day 1, the 20 and 40 mg/mL groups had significantly lower nicotine levels at 120 minutes relative to 0 minutes post-nicotine exposure (p=0.053 and p=0.013, respectively). The 80 mg/mL group exhibited lower nicotine levels at 60 minutes relative to 0 minutes post-nicotine exposure (p=0.005), but not at 120 minutes (p=0.194), presumably due to higher data variability at that time point. On day 10, only the 80 mg/mL group had significantly lower nicotine levels at 120 minutes relative to 0 minutes post-nicotine (p=0.022) (Figure 2b).

**Fig. 2.**
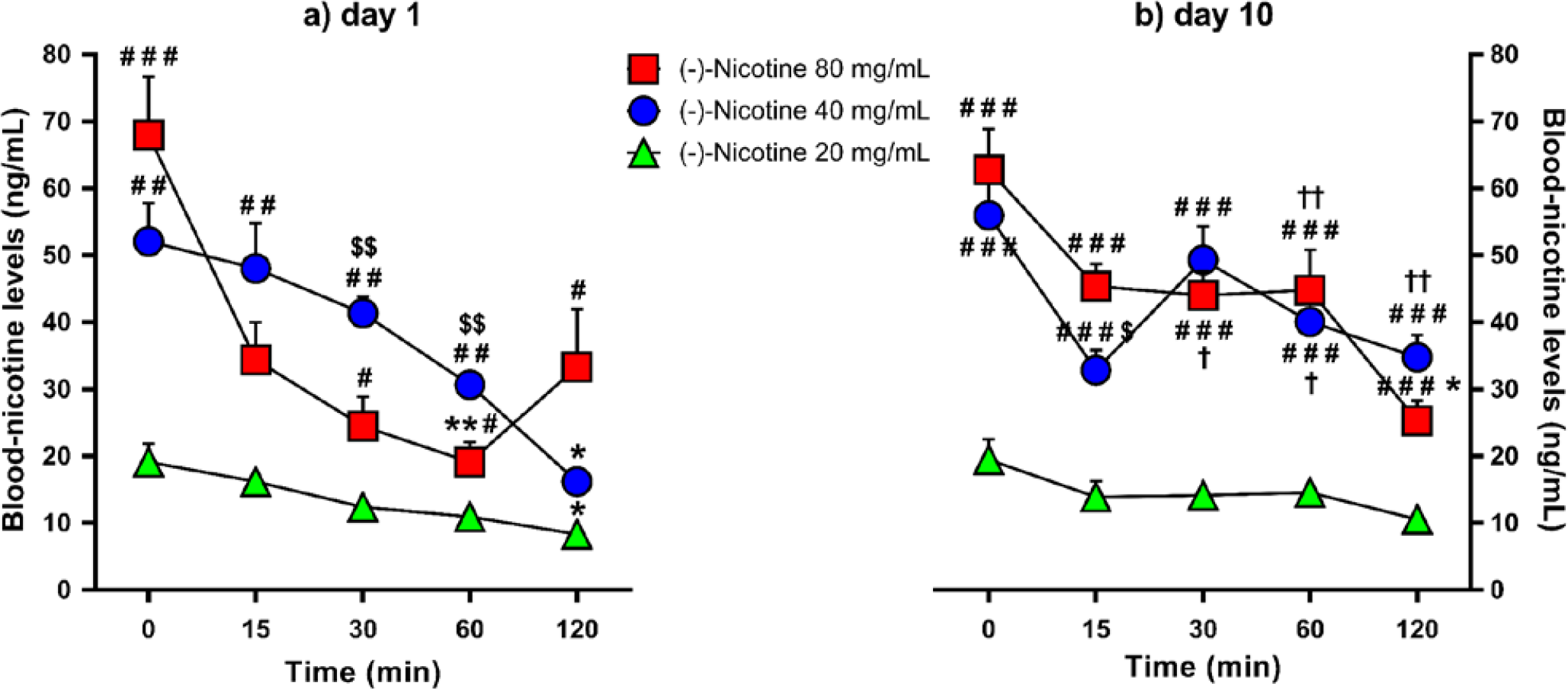
Mean blood-nicotine levels (+SEM) over time in the 20, 40 and 80 mg/mL nicotine concentration groups (N=6/group) following the 1^st^ (**panel a**) and 10^th^ (**panel b**) day of nicotine e-vape exposure. * p≤0.05 and ** p≤0.01 compared with baseline; # p≤0.05, ## p≤0.01 and ### p≤0.001 compared with the 20 mg/mL nicotine group; $ p≤0.05 and $$ p≤0.01 compared with the 80 mg/mL nicotine group; † p≤0.05 and †† p≤0.01 compared with day 1. Significance symbols are reported over and under the corresponding dataset.

On days 1 and 10, the 40 and 80 mg/mL nicotine groups had higher nicotine levels than the 20 mg/mL group at all the time points (Figures 2a & 2b), except on day 1 at 15 (for *20 vs*. *80 mg*/*mL* comparison) and 120 minutes (for *20 vs*. *40 mg*/*mL* comparisons) post-nicotine exposure (*20 vs*. *40 mg*/*mL*: day 1, p≤0.01 for all comparisons except the 120-min time point; day 10, p<0.001 for all comparisons; *20 vs*. *80 mg*/*mL*: day 1, p≤0.001 for 0 minutes, and p≤0.05 for all comparisons except for the 15-min time point; day 10, p≤0.001 for all comparisons). Relative to the 40 mg/mL nicotine group, rats exposed to 80 mg/mL nicotine exhibited lower blood-nicotine levels on day 1 at the 30- and 60-min time points (p<0.01 for all comparisons) but higher nicotine levels on day 10 at the 15-min time point (p=0.025; Figures 2a & 2b).

Analysis of blood-nicotine levels over days (day 1 vs. day 10) indicated higher blood-nicotine levels on day 10 at the 60-min and 120-min time points for the 40 mg/mL nicotine group (p=0.019 and p=0.002, respectively), and at the 30-min and 60-min time points for the 80 mg/mL nicotine group (p=0.020 and p=0.007, respectively; Figures 2a & 2b).

### Plasma-cotinine levels

Analysis of plasma-cotinine levels yielded significant main effects of day [F(1,15) =12.373, p=0.003], time [F(8,60)=138.612, p<0.001] and group [F(2,15)=43.703, p<0.001). There were also significant day × group interaction effect [F(2,15)=7.585, p=0.005] and a significant time × group interaction effect [F(4,60)=138.612, p=0.001]. There were not significant day × time [F(4,60)=1.41, p=0.239] or day × time × group [F(8,60)=0.761, p=0.638] interaction effects. As shown in Figures 3a & 3b, Bonferroni post-hoc comparisons confirmed that all the three nicotine concentration groups exhibited higher blood-cotinine levels at 120 minutes relative to 0 minutes post-nicotine on days 1 and 10 (*20 and 40 mg*/*mL*, p<0.001; *80 mg*/*mL*, p=0.003). Furthermore, the 40 and 80 mg/mL nicotine groups exhibited higher blood-cotinine levels over time relative to the 20 mg/mL group on day 1 (Figure 3a: *20 vs*. *40 mg*/*mL*: p=0.016; *20 vs*. *80 mg*/*mL*: p=0.002) and day 10 (Figure 3b: p<0.001 for all comparisons), while the 80 mg/mL nicotine group exhibited higher blood-cotinine levels relative to the 40 mg/mL group on day 10 (Figure 3b: p<0.001). The 80 mg/mL nicotine group also exhibited higher blood-cotinine levels on day 10 relative to day 1 across all time points (p=0.006) (Figures 3a & 3b).

**Fig. 3.**
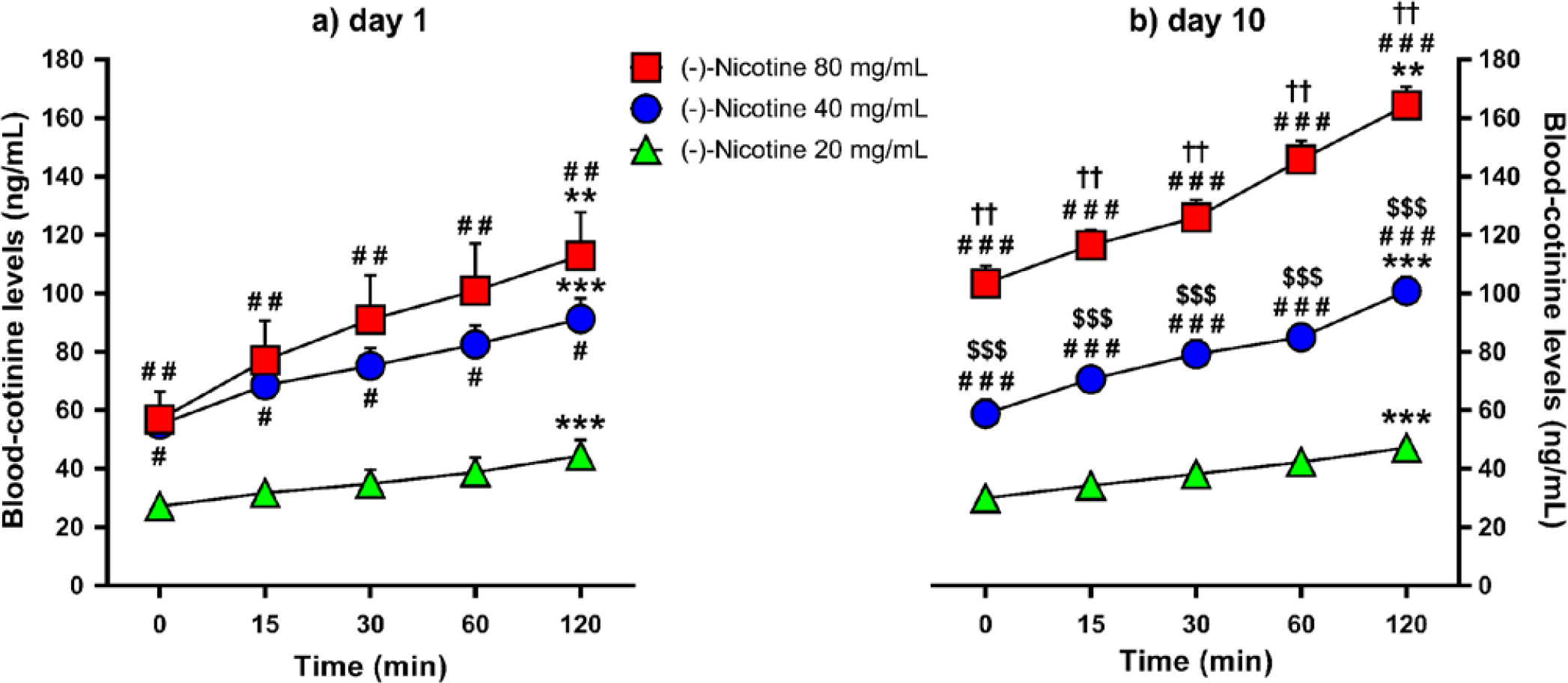
Mean blood-cotinine levels (+SEM) over time in the 20, 40 and 80 mg/mL nicotine concentration groups (N=6/group) following the 1^st^ (**panel a**) and 10^th^ (**panel b**) day of nicotine e-vape exposure. ** p≤0.01 and **** p≤0.001 compared with baseline; # p≤0.05, ## p≤0.01 and ### p≤0.001 compared with the 20 mg/mL nicotine group; $$$ p≤0.001 compared with the 80 mg/mL nicotine group; †† p≤0.01 compared with day 1. Significance symbols are reported over and under the corresponding dataset.

### Spontaneous withdrawal symptoms 24 hours after single or repeated nicotine exposure

Analysis of spontaneous physical withdrawal symptoms 24 hours into nicotine withdrawal indicated a significant main effect of session [F(3,45)=75.740, p<0.001] but not nicotine concentration group [F(2,15)=0.775, p=0.478]. There was also a significant session × group interaction effect on spontaneous withdrawal scores [F(6,45)=10.498, p<0.001] (Figure 4). Post-hoc comparisons indicated that the 20 and 40 mg/mL group exhibited higher withdrawal scores on day 1, 10 and 11 relative to baseline (*20 mg*/*mL*: p≤0.05 on day 1 and 11, p=0.009 on day 10; *40 mg*/*mL*: p=0.015 on day 1, p≤0.001 on day 10 and 11). The 40 mg/mL group exhibited also a higher withdrawal score on day 11 (precipitated withdrawal) relative to spontaneous withdrawal on day 1 and 10 (p≤0.001 and p=0.016, respectively). The 80 mg/mL exhibited higher withdrawal scores on days 10 and 11 relative to baseline (p=0.001 and p=0.035, respectively) and day 1 (p<0.001 and p=0.050, respectively). This different pattern of results across nicotine concentrations may be due in part to the chosen WD time point (24 hours).

**Fig. 4.**
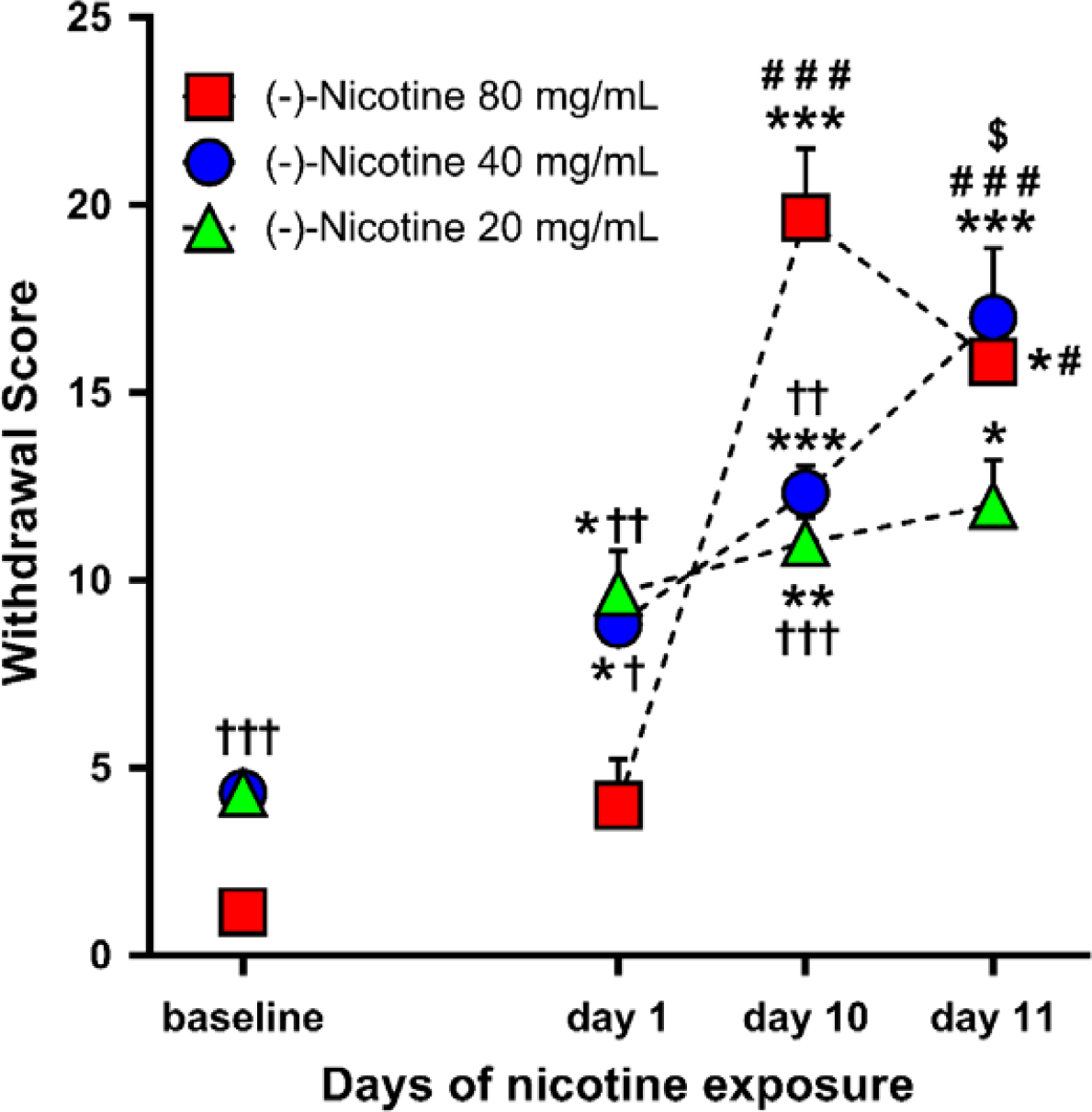
Mean withdrawal score (+SEM) in baseline and 24 hours post-nicotine e-vape exposure on day 1 and 10 (spontaneous withdrawal), and day 11 (precipitated withdrawal) for the 20, 40 and 80 mg/mL nicotine concentration groups (N=6/group) * p≤0.05, ** p≤0.01 and *** p≤0.001 compared with baseline; # p≤0.05 and ### p≤0.001 compared with day 1; $ p≤0.05 compared with day 10; † p≤0.05, †† p≤0.01 and ††† p≤0.001 compared with 80 mg/mL nicotine concentration group. Significance symbols are reported over, beside and under the corresponding dataset.

As a result of random group assignment, the 80 mg/mL group exhibited lower baseline (i.e., pre-nicotine) “withdrawal scores” in comparison to the 20 and 40 mg/mL nicotine groups (p≤0.001 for both comparisons). After the first 1-hour nicotine session, the 80 mg/mL group again exhibited lower withdrawal scores in comparison to the 20 and 40 mg/mL nicotine groups (p=0.004 and p=0.012, respectively). After the 10^th^ 1-hour nicotine session, the 80 mg/mL group exhibited *higher* withdrawal scores in comparison to the 20 and 40 mg/mL nicotine groups (p<0.001 and p=0.002, respectively). After the 11^th^ 1-hour nicotine session and mecamylamine injection, there were no significant group differences in withdrawal scores.

### Spontaneous withdrawal symptoms after 10 days of nicotine vapor inhalation

A 2-way ANOVA of baseline and day 10 spontaneous withdrawal symptoms indicated a significant main effect of time [F(5,75)=3.756, p<0.001] and group [F(2,15)=20.943, p<0.001] and a significant time × group interaction [F(10,75)=7.512, p<0.001]. As shown in Figure 5, after 10 days of nicotine vapor inhalation, the 20 mg/mL group exhibited a higher withdrawal score relative to baseline only at 24 hours post-nicotine exposure (p=0.022), while the 40 and 80 mg/mL groups exhibited higher withdrawal scores relative to baseline at all the time points (*40 mg*/*mL*, p<0.05 at 1 and 6 hours, p=0.002 at 24 hours, and p<0.001 at 2 and 4 hours post-nicotine; *80 mg*/*mL*, p<0.01 for all comparisons). The 80 mg/mL nicotine group exhibited also higher withdrawal scores at 4-24 hours relative to 1-2 hours post-nicotine exposure (p<0.05 for all comparisons).

**Fig. 5.**
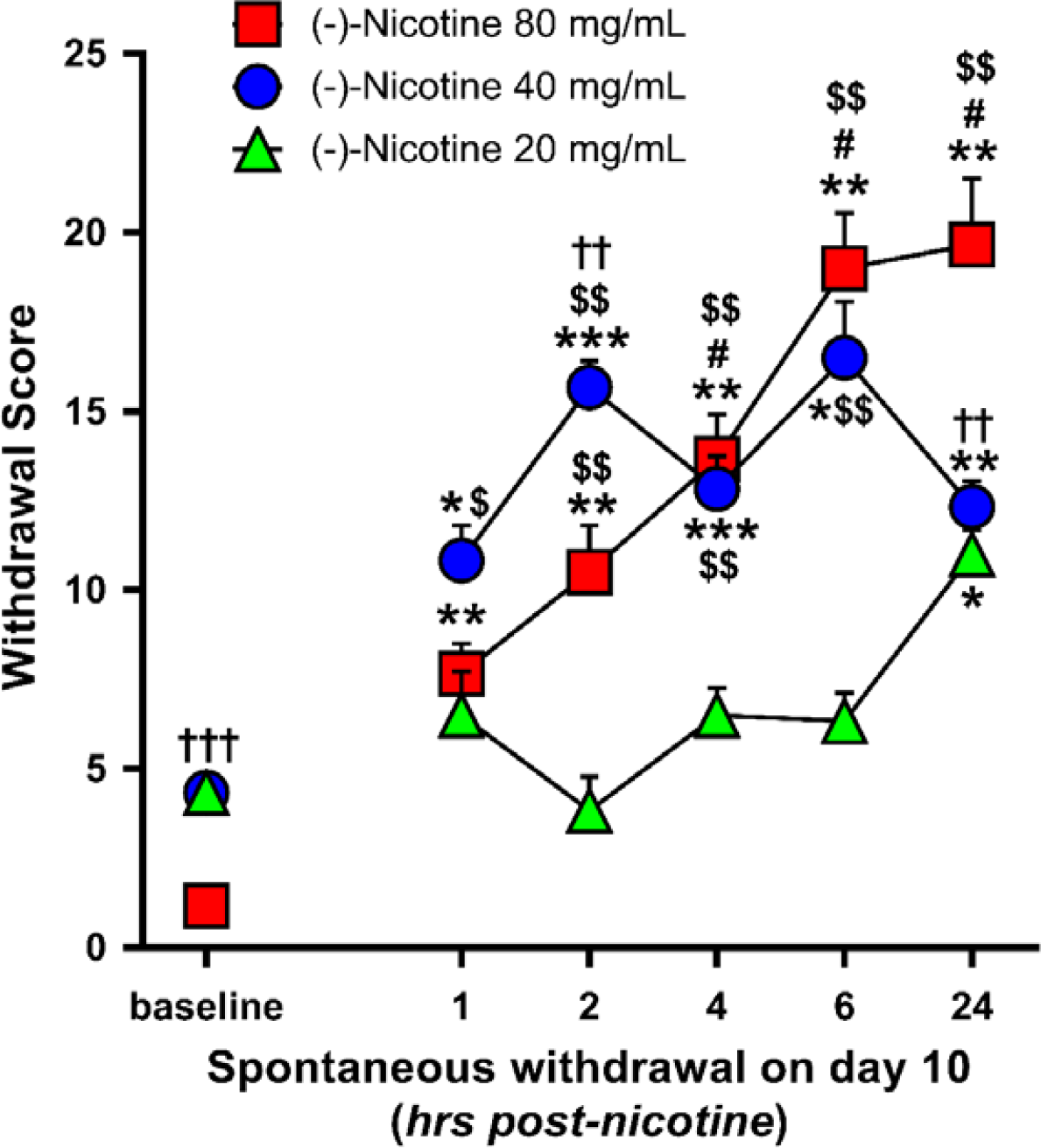
Mean spontaneous withdrawal scores (+SEM) at 1, 2, 4, 6 & 24 hours post-nicotine exposure following the 10^th^ day of nicotine e-vape exposure for the 20, 40 and 80 mg/mL nicotine concentration groups (N=6/group). * p≤0.05, ** p≤0.01 and *** p≤0.001 compared with baseline; # p≤0.05 compared with 1 and 2 hours post-nicotine; $ p≤0.05 and $$ p≤0.01 compared with the 20 mg/mL nicotine concentration group; †† p≤0.01 and ††† p≤0.001 compared with the 80 mg/mL nicotine concentration group. Significance symbols are reported over and under the corresponding dataset.

As follow-up to the main effect of nicotine concentration, the 40 and 80 mg/mL groups exhibited higher withdrawal scores over time than the 20 mg/mL group (p<0.001 for both comparisons). More specifically, the 40 mg/mL nicotine group exhibited higher withdrawal scores than the 20 mg/mL nicotine group at 1, 2, 4 and 6 hours post-nicotine (p<0.05 at 1 hour, and p≤0.001 for the other comparisons), while the 80 mg/mL nicotine group exhibited higher withdrawal scores than the 20 mg/mL nicotine group at 2, 4, 6 and 24 hours post-nicotine (p≤0.001 for all comparisons). The 40 mg/mL nicotine group exhibited higher withdrawal scores than the 80 mg/mL nicotine group at 2 hours post-nicotine (p=0.009), whereas the 80 mg/mL nicotine group exhibited higher withdrawal scores than the 40 mg/mL nicotine group at 24 hours post-nicotine (p=0.002).

### Precipitated withdrawal symptoms after 11 days of nicotine vapor inhalation

A 2-way ANOVA of baseline and day 11 precipitated withdrawal symptoms indicated a significant main effect of time [F(6,90) = 41.264, p<0.001] and group [F(2,15) = 6.776, p=0.008] and a significant time × group interaction [F(12,90)=7.829, p<0.001). As shown in Figure 6, after 11 days of nicotine vapor inhalation all the three nicotine concentration groups exhibited higher precipitated withdrawal scores relative to baseline: the 20 mg/mL group at 1, 2, 4, 24 and 48 hours post-nicotine (p≤0.001 at 1 and 48 hours, and ≤0.05 for the other comparisons), the 40 mg/mL group at all the post-nicotine time points (p<0.05 at 1 and 2 hours, p<0.01 at 4 and 6 hours, and p≤0.001 at 24 and 48 hours), and the 80 mg/mL group at 1, 2, 4, 6 and 48 hours post-nicotine (p≤0.01 for all comparisons except at 1 hour post-nicotine, p=0.042). The 20 mg/mL group exhibited also lower withdrawal scores at 4, 6 and 24 hours relative to 1 hour post-nicotine (p=0.005 at 4 hours, and p<0.05 at 6 and 24 hours), and a higher withdrawal score at 48 hours relative to 4 and 6 hours post-nicotine (p<0.01 for all comparisons). We also observed a higher withdrawal score at 48 hours relative to 6 hours post-nicotine exposure in the 40 mg/mL group (p=0.025).

**Fig. 6.**
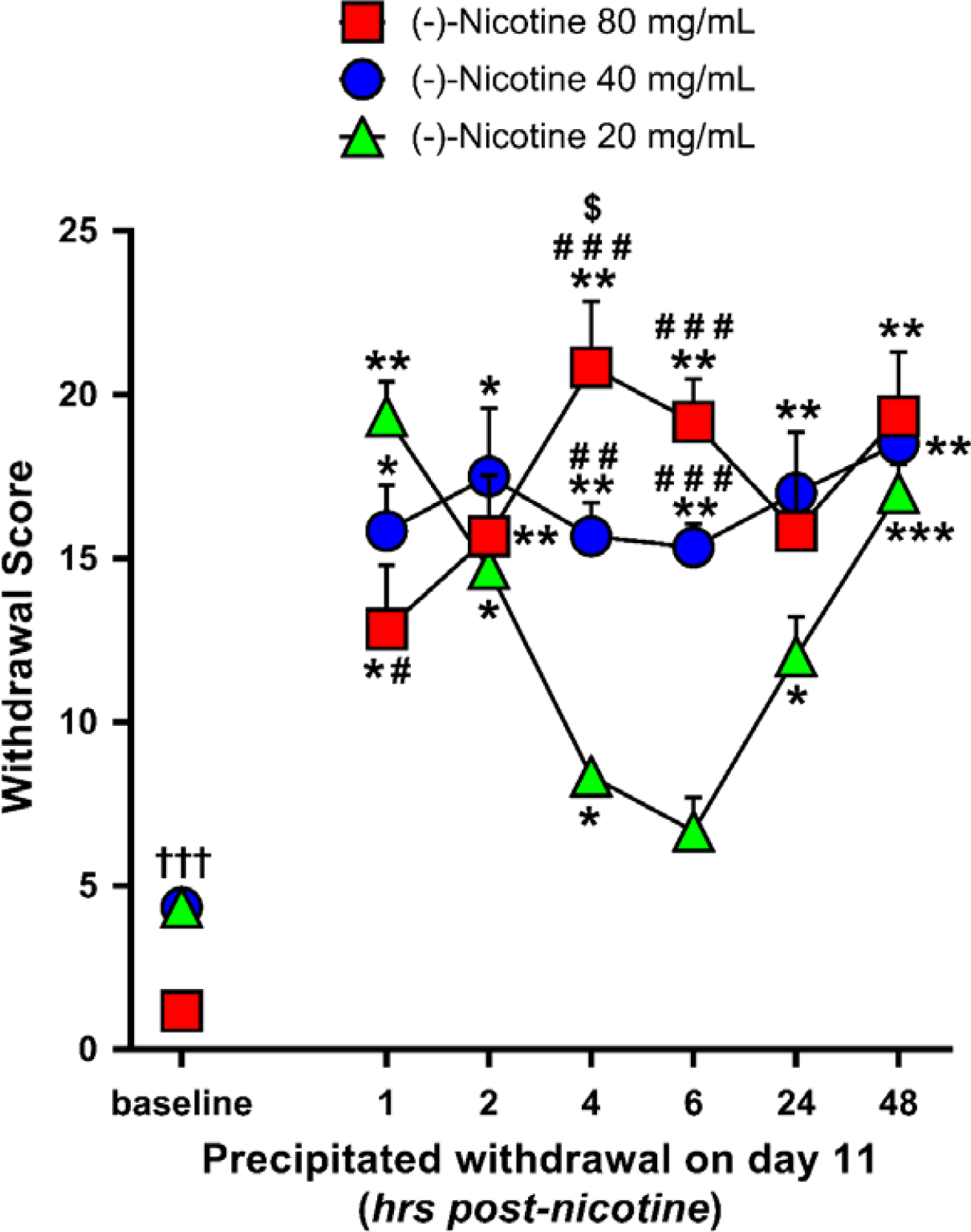
Mean precipitated withdrawal scores (+SEM) at 1, 2, 4, 6, 24 & 48 hours post-nicotine exposure on the 11^th^ day of nicotine e-vape exposure for the 20, 40 and 80 mg/mL nicotine concentration groups (N=6/group); * p≤0.05, ** p≤0.01 and *** p≤0.001 compared with baseline; # p≤0.05, # # p≤0.01 and # # # p≤0.001 compared with the 20 mg/mL nicotine concentration group; $ p≤0.05 compared with the 40 mg/mL nicotine concentration group. Significance symbols are reported over, beside and under the corresponding dataset.

As follow-up to the main effect of nicotine concentration, 40 and 80 mg/mL groups exhibited higher withdrawal scores in comparison to the 20 mg/mL group (p≤0.05 for all comparisons). Specifically, the 40 and 80 mg/mL nicotine groups exhibited higher withdrawal scores than the 20 mg/mL group at 4 and 6 hours post-nicotine exposure (*20 vs*. *40 mg*/*mL*: p=0.004 at 4 hours, and p<0.001 at 6 hours; *20 vs*. *80 mg*/*mL*: p<0.001 for all comparisons). Moreover, the 20 mg/mL group exhibited a higher withdrawal score relative to the 80 mg/mL group at 1 hour post-nicotine exposure (p=0.025), while the 80 mg/mL nicotine group exhibited a higher withdrawal score relative to the 40 mg/mL group at 4 hours post-nicotine exposure (p=0.044).

Within each nicotine concentration group, paired t-tests (2-tails) between spontaneous (on day 10) and precipitated (day 11) withdrawal indicated higher *precipitated* withdrawal scores at 1 and 2 hours post-nicotine exposure for the 20 mg/mL nicotine group (p<0.001 and p=0.002, respectively), at 1 and 24 hours post-nicotine exposure for the 40 mg/mL group (p=0.023 and p=0.003, respectively), and at 2 and 4 hours post-nicotine exposure for the 80 mg/mL group (p=0.001 and p=0.004, respectively).

Tables 1 and 2 contain data for individual somatic signs of nicotine withdrawal over time on the days 1, 10 and 11. Table 1 shows scores for individual somatic withdrawal signs during spontaneous withdrawal testing. Table 2 shows scores for individual somatic withdrawal signs during mecamylamine-precipitated withdrawal testing. For both spontaneous and precipitated withdrawal tests and across nicotine concentration groups, the most frequent symptom categories were grooming/licking, scratch/bite and body shakes.

**Table 1.**
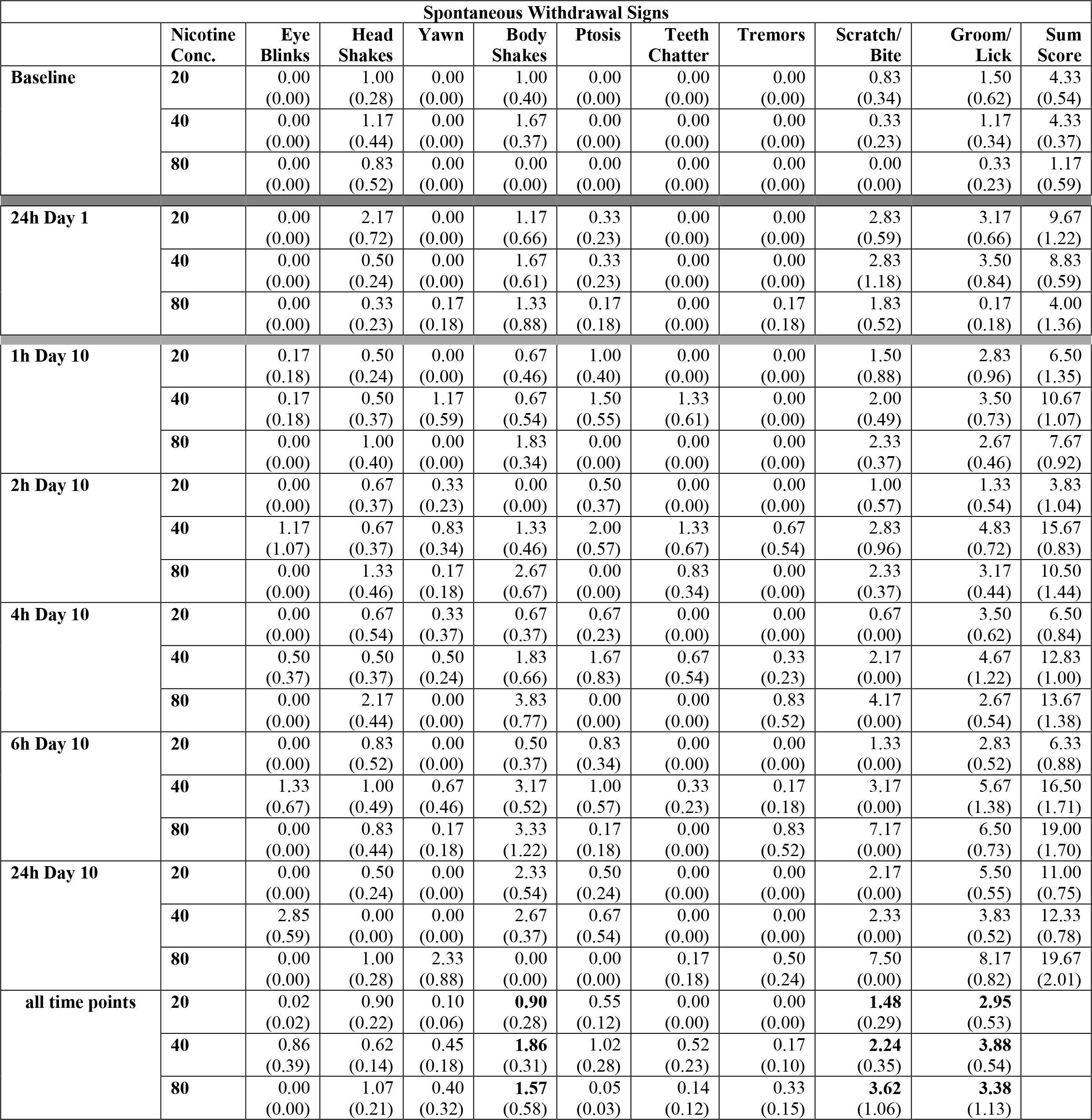
Mean number (+SEM) of individual somatic signs in baseline and *spontaneous withdrawal* on day 1 (at 24 hours post-nicotine exposure) and day 10 (at 1, 2, 4, 6 & 24 hours post-nicotine exposure) for the 20, 40 and 80 mg/mL nicotine concentration groups (N=6/group). *The “all time points” row indicates the mean number (+SEM) of each symptom averaged across all the time points.*

| Spontaneous Withdrawal Signs |  |  |  |  |  |  |  |  |  |  |  |
| --- | --- | --- | --- | --- | --- | --- | --- | --- | --- | --- | --- |
|  | Nicotine Conc. | Eye Blinks | Head Shakes | Yawn | Body Shakes | Ptois | Teeth Chatter | Tremors | Scratch/ Bite | Groom/ Lick | Sum Score |
| Baseline | 20 | 0.00<br>(0.00) | 1.00<br>(0.28) | 0.00<br>(0.00) | 1.00<br>(0.40) | 0.00<br>(0.00) | 0.00<br>(0.00) | 0.00<br>(0.00) | 0.83<br>(0.34) | 1.50<br>(0.62) | 4.33<br>(0.54) |
|  | 40 | 0.00<br>(0.00) | 1.17<br>(0.44) | 0.00<br>(0.00) | 1.67<br>(0.37) | 0.00<br>(0.00) | 0.00<br>(0.00) | 0.00<br>(0.00) | 0.33<br>(0.23) | 1.17<br>(0.34) | 4.33<br>(0.37) |
|  | 80 | 0.00<br>(0.00) | 0.83<br>(0.52) | 0.00<br>(0.00) | 0.00<br>(0.00) | 0.00<br>(0.00) | 0.00<br>(0.00) | 0.00<br>(0.00) | 0.00<br>(0.00) | 0.33<br>(0.23) | 1.17<br>(0.59) |
| 24h Day 1 | 20 | 0.00<br>(0.00) | 2.17<br>(0.72) | 0.00<br>(0.00) | 1.17<br>(0.66) | 0.33<br>(0.23) | 0.00<br>(0.00) | 0.00<br>(0.00) | 2.83<br>(0.59) | 3.17<br>(0.66) | 9.67<br>(1.22) |
|  | 40 | 0.00<br>(0.00) | 0.50<br>(0.24) | 0.00<br>(0.00) | 1.67<br>(0.61) | 0.33<br>(0.23) | 0.00<br>(0.00) | 0.00<br>(0.00) | 2.83<br>(1.18) | 3.50<br>(0.84) | 8.83<br>(0.59) |
|  | 80 | 0.00<br>(0.00) | 0.33<br>(0.23) | 0.17<br>(0.18) | 1.33<br>(0.88) | 0.17<br>(0.18) | 0.00<br>(0.00) | 0.17<br>(0.18) | 1.83<br>(0.52) | 0.17<br>(0.18) | 4.00<br>(1.36) |
| 1h Day 10 | 20 | 0.17<br>(0.18) | 0.50<br>(0.24) | 0.00<br>(0.00) | 0.67<br>(0.46) | 1.00<br>(0.40) | 0.00<br>(0.00) | 0.00<br>(0.00) | 1.50<br>(0.88) | 2.83<br>(0.96) | 6.50<br>(1.35) |
|  | 40 | 0.17<br>(0.18) | 0.50<br>(0.37) | 1.17<br>(0.59) | 0.67<br>(0.54) | 1.50<br>(0.55) | 1.33<br>(0.61) | 0.00<br>(0.00) | 2.00<br>(0.49) | 3.50<br>(0.73) | 10.67<br>(1.07) |
|  | 80 | 0.00<br>(0.00) | 1.00<br>(0.40) | 0.00<br>(0.00) | 1.83<br>(0.34) | 0.00<br>(0.00) | 0.00<br>(0.00) | 0.00<br>(0.00) | 2.33<br>(0.37) | 2.67<br>(0.46) | 7.67<br>(0.92) |
| 2h Day 10 | 20 | 0.00<br>(0.00) | 0.67<br>(0.37) | 0.33<br>(0.23) | 0.00<br>(0.00) | 0.50<br>(0.37) | 0.00<br>(0.00) | 0.00<br>(0.00) | 1.00<br>(0.57) | 1.33<br>(0.54) | 3.83<br>(1.04) |
|  | 40 | 1.17<br>(1.07) | 0.67<br>(0.37) | 0.83<br>(0.34) | 1.33<br>(0.46) | 2.00<br>(0.57) | 1.33<br>(0.67) | 0.67<br>(0.54) | 2.83<br>(0.96) | 4.83<br>(0.72) | 15.67<br>(0.83) |
|  | 80 | 0.00<br>(0.00) | 1.33<br>(0.46) | 0.17<br>(0.18) | 2.67<br>(0.67) | 0.00<br>(0.00) | 0.83<br>(0.34) | 0.00<br>(0.00) | 2.33<br>(0.37) | 3.17<br>(0.44) | 10.50<br>(1.44) |
| 4h Day 10 | 20 | 0.00<br>(0.00) | 0.67<br>(0.54) | 0.33<br>(0.37) | 0.67<br>(0.37) | 0.67<br>(0.23) | 0.00<br>(0.00) | 0.00<br>(0.00) | 0.67<br>(0.00) | 3.50<br>(0.62) | 6.50<br>(0.84) |
|  | 40 | 0.50<br>(0.37) | 0.50<br>(0.37) | 0.50<br>(0.24) | 1.83<br>(0.66) | 1.67<br>(0.83) | 0.67<br>(0.54) | 0.33<br>(0.23) | 2.17<br>(0.00) | 4.67<br>(1.22) | 12.83<br>(1.00) |
|  | 80 | 0.00<br>(0.00) | 2.17<br>(0.44) | 0.00<br>(0.00) | 3.83<br>(0.77) | 0.00<br>(0.00) | 0.00<br>(0.00) | 0.83<br>(0.52) | 4.17<br>(0.00) | 2.67<br>(0.54) | 13.67<br>(1.38) |
| 6h Day 10 | 20 | 0.00<br>(0.00) | 0.83<br>(0.52) | 0.00<br>(0.00) | 0.50<br>(0.37) | 0.83<br>(0.34) | 0.00<br>(0.00) | 0.00<br>(0.00) | 1.33<br>(0.00) | 2.83<br>(0.52) | 6.33<br>(0.88) |
|  | 40 | 1.33<br>(0.67) | 1.00<br>(0.49) | 0.67<br>(0.46) | 3.17<br>(0.52) | 1.00<br>(0.57) | 0.33<br>(0.23) | 0.17<br>(0.18) | 3.17<br>(0.00) | 5.67<br>(1.38) | 16.50<br>(1.71) |
|  | 80 | 0.00<br>(0.00) | 0.83<br>(0.44) | 0.17<br>(0.18) | 3.33<br>(1.22) | 0.17<br>(0.18) | 0.00<br>(0.00) | 0.83<br>(0.52) | 7.17<br>(0.00) | 6.50<br>(0.73) | 19.00<br>(1.70) |
| 24h Day 10 | 20 | 0.00<br>(0.00) | 0.50<br>(0.24) | 0.00<br>(0.00) | 2.33<br>(0.54) | 0.50<br>(0.24) | 0.00<br>(0.00) | 0.00<br>(0.00) | 2.17<br>(0.00) | 5.50<br>(0.55) | 11.00<br>(0.75) |
|  | 40 | 2.85<br>(0.59) | 0.00<br>(0.00) | 0.00<br>(0.00) | 2.67<br>(0.37) | 0.67<br>(0.54) | 0.00<br>(0.00) | 0.00<br>(0.00) | 2.33<br>(0.00) | 3.83<br>(0.52) | 12.33<br>(0.78) |
|  | 80 | 0.00<br>(0.00) | 1.00<br>(0.28) | 2.33<br>(0.88) | 0.00<br>(0.00) | 0.00<br>(0.00) | 0.17<br>(0.18) | 0.50<br>(0.24) | 7.50<br>(0.00) | 8.17<br>(0.82) | 19.67<br>(2.01) |
| all time points | 20 | 0.02<br>(0.02) | 0.90<br>(0.22) | 0.10<br>(0.06) | **0.90**<br>(0.28) | 0.55<br>(0.12) | 0.00<br>(0.00) | 0.00<br>(0.00) | **1.48**<br>(0.29) | **2.95**<br>(0.53) |  |
|  | 40 | 0.86<br>(0.39) | 0.62<br>(0.14) | 0.45<br>(0.18) | **1.86**<br>(0.31) | 1.02<br>(0.28) | 0.52<br>(0.23) | 0.17<br>(0.10) | **2.24**<br>(0.35) | **3.88**<br>(0.54) |  |
|  | 80 | 0.00<br>(0.00) | 1.07<br>(0.21) | 0.40<br>(0.32) | **1.57**<br>(0.58) | 0.05<br>(0.03) | 0.14<br>(0.12) | 0.33<br>(0.15) | **3.62**<br>(1.06) | **3.38**<br>(1.13) |  |

**Table 2.** Mean number (+SEM) of individual somatic signs on day 11 (*precipitated withdrawal*) at 1, 2, 4, 6, 24 & 48 hours post-nicotine exposure for the 20, 40 and 80 mg/mL nicotine concentration groups (N=6/group). *The “all time points” row indicates the mean number (+SEM) of each symptom averaged across all the time points*

| Mecamylamine-Precipitated Withdrawal Signs |  |  |  |  |  |  |  |  |  |  |  |
| --- | --- | --- | --- | --- | --- | --- | --- | --- | --- | --- | --- |
|  | Nicotine Conc. | Eye Blinks | Head Shakes | Yawn | Body Shakes | Ptosis | Teeth Chatter | Tremors | Scratch/ Bite | Grooming/ Licking | Sum Score |
| **1h Day 11** | **20** | 0.50<br>(0.37) | 0.50<br>(0.55) | 0.17<br>(0.18) | 3.00<br>(0.75) | 1.17<br>(0.34) | 0.17<br>(0.18) | 0.83<br>(0.34) | 5.33<br>(0.92) | 7.67<br>(0.61) | 19.33<br>(1.15) |
|  | **40** | 0.67<br>(0.46) | 0.67<br>(0.37) | 0.17<br>(0.18) | 4.17<br>(0.66) | 1.00<br>(0.40) | 0.17<br>(0.18) | 0.17<br>(0.18) | 3.33<br>(0.83) | 5.50<br>(0.24) | 15.83<br>(1.53) |
|  | **80** | 0.83<br>(0.44) | 1.33<br>(0.54) | 0.00<br>(0.00) | 2.50<br>(1.52) | 1.33<br>(0.46) | 0.00<br>(0.00) | 0.33<br>(0.23) | 0.50<br>(0.37) | 6.00<br>(0.75) | 12.83<br>(2.14) |
| **2h Day 11** | **20** | 0.83<br>(0.34) | 0.67<br>(0.23) | 0.17<br>(0.18) | 2.00<br>(0.63) | 1.83<br>(0.44) | 0.17<br>(0.18) | 1.00<br>(0.57) | 3.33<br>(0.92) | 4.67<br>(0.88) | 14.67<br>(1.29) |
|  | **40** | 1.00<br>(0.40) | 2.17<br>(0.82) | 1.00<br>(0.57) | 2.33<br>(0.67) | 1.50<br>(0.37) | 0.00<br>(0.00) | 0.50<br>(0.37) | 5.00<br>(1.30) | 4.00<br>(1.13) | 17.50<br>(2.28) |
|  | **80** | 0.50<br>(0.24) | 1.50<br>(0.68) | 0.17<br>(0.18) | 2.50<br>(0.68) | 1.17<br>(0.44) | 0.00<br>(0.00) | 0.00<br>(0.00) | 5.00<br>(0.85) | 4.83<br>(0.87) | 15.67<br>(2.03) |
| **4h Day 11** | **20** | 0.33<br>(0.23) | 0.17<br>(0.18) | 0.17<br>(0.18) | 1.67<br>(0.37) | 1.50<br>(0.37) | 0.00<br>(0.00) | 0.67<br>(0.37) | 2.17<br>(0.77) | 1.67<br>(0.61) | 8.33<br>(0.37) |
|  | **40** | 0.50<br>(0.55) | 0.17<br>(0.18) | 0.33<br>(0.23) | 3.50<br>(0.37) | 0.50<br>(0.37) | 0.00<br>(0.00) | 0.00<br>(0.00) | 5.00<br>(0.69) | 5.67<br>(0.88) | 15.67<br>(1.12) |
|  | **80** | 0.33<br>(0.37) | 2.83<br>(0.52) | 0.67<br>(0.37) | 1.83<br>(0.66) | 0.67<br>(0.73) | 0.00<br>(0.00) | 0.00<br>(0.00) | 7.33<br>(1.35) | 7.17<br>(0.91) | 20.83<br>(2.22) |
| **6h Day 11** | **20** | 0.33<br>(0.37) | 0.50<br>(0.37) | 0.67<br>(0.23) | 1.00<br>(0.28) | 1.00<br>(0.28) | 0.00<br>(0.00) | 0.67<br>(0.37) | 0.83<br>(0.34) | 1.67<br>(0.83) | 6.67<br>(1.12) |
|  | **40** | 0.17<br>(0.18) | 1.50<br>(0.79) | 0.00<br>(0.00) | 3.67<br>(0.73) | 1.00<br>(0.57) | 0.00<br>(0.00) | 0.17<br>(0.18) | 3.50<br>(0.47) | 5.33<br>(0.92) | 15.33<br>(0.78) |
|  | **80** | 0.67<br>(0.37) | 1.00<br>(0.57) | 1.67<br>(0.37) | 0.33<br>(0.23) | 1.00<br>(0.40) | 0.17<br>(0.18) | 0.67<br>(0.54) | 4.33<br>(1.51) | 9.33<br>(1.12) | 19.17<br>(1.43) |
| **24h Day 11** | **20** | 0.83<br>(0.34) | 0.00<br>(0.00) | 0.17<br>(0.18) | 1.83<br>(0.34) | 1.17<br>(0.44) | 0.00<br>(0.00) | 0.00<br>(0.00) | 2.33<br>(0.67) | 5.67<br>(0.67) | 12.00<br>(1.33) |
|  | **40** | 0.00<br>(0.00) | 3.00<br>(0.80) | 0.50<br>(0.24) | 1.83<br>(0.59) | 1.67<br>(1.08) | 0.00<br>(0.00) | 0.00<br>(0.00) | 5.33<br>(0.46) | 4.67<br>(0.46) | 17.00<br>(0.49) |
|  | **80** | 0.00<br>(0.00) | 1.83<br>(0.72) | 0.17<br>(0.18) | 1.67<br>(1.15) | 0.33<br>(0.23) | 0.00<br>(0.00) | 0.17<br>(0.18) | 4.67<br>(1.01) | 7.00<br>(2.48) | 15.83<br>(3.30) |
| **48h Day 11** | **20** | 1.50<br>(0.47) | 1.20<br>(0.42) | 0.00<br>(0.00) | 3.50<br>(0.97) | 0.67<br>(0.23) | 0.17<br>(0.18) | 0.00<br>(0.00) | 4.00<br>(0.69) | 6.17<br>(0.87) | 17.00<br>(0.94) |
|  | **40** | 1.33<br>(0.61) | 0.83<br>(0.72) | 0.00<br>(0.00) | 3.50<br>(1.22) | 1.67<br>(0.46) | 0.00<br>(0.00) | 0.50<br>(0.55) | 5.00<br>(0.75) | 5.67<br>(1.08) | 18.50<br>(1.09) |
|  | **80** | 0.33<br>(0.37) | 2.17<br>(0.96) | 0.00<br>(0.00) | 3.83<br>(1.04) | 1.00<br>(0.28) | 0.00<br>(0.00) | 0.00<br>(0.00) | 5.83<br>(0.96) | 6.17<br>(1.84) | 19.33<br>(2.15) |
| **all time points** | **20** | 0.72<br>(0.17) | 0.51<br>(0.16) | 0.23<br>(0.09) | **2.17**<br>(0.35) | 1.22<br>(0.15) | 0.09<br>(0.04) | 0.53<br>(0.16) | **3.00**<br>(0.59) | **4.59**<br>(0.93) |  |
|  | **40** | 0.61<br>(0.19) | 1.39<br>(0.40) | 0.33<br>(0.14) | **3.17**<br>(0.34) | 1.22<br>(0.18) | 0.03<br>(0.03) | 0.22<br>(0.09) | **4.53**<br>(0.33) | **5.14**<br>(0.25) |  |
|  | **80** | 0.25<br>(0.37) | 1.78<br>(0.25) | 0.45<br>(0.24) | **2.11**<br>(0.44) | 0.92<br>(0.14) | 0.03<br>(0.08) | 0.20<br>(0.10) | **4.61**<br>(0.86) | **6.75**<br>(0.57) |  |

## Discussion

Exposing male Wistar rats to non-contingent nicotine vapor inhalation using ENDS produces somatic withdrawal symptoms and measurable blood-nicotine and blood-cotinine levels that change according to 1) concentration of nicotine in vape solution, 2) number of days of nicotine vapor exposure, 3) time since termination of nicotine vapor exposure, and 4) relative to the withdrawal signs, whether withdrawal was spontaneous or precipitated (by systemic injection of mecamylamine). This is the first concentration-dependent time course analysis of somatic withdrawal signs and blood-nicotine/cotinine levels in rats after single or repeated bouts of nicotine vapor inhalation using ENDS resembling those currently used by humans.

ENDS are relatively new for the field and are being used by multiple labs to vaporize and deliver various drugs to rodents via inhalation. Published work shows that ENDS can be used to vaporize and deliver nicotine (Javadi-Paydar et al. 2019), THC (Nguyen et al. 2016a, 2018; Javadi-Paydar et al. 2018, 2019), opiates (Vendruscolo et al. 2018), stimulants (Nguyen et al. 2017), or combinations of these drugs (Javadi-Paydar et al. 2019) to rats on experimenter-determined schedules, and that these exposures produce expected blood-drug levels, relative to prior work using other systemic routes of drug administration. Furthermore, for many of these drugs, labs are working to establish reliable e-cigarette vapor self-administration using operant tasks that can be learned, extinguished, and altered according to drug concentrations in the vape solution. Not surprisingly, this has been easier for some drugs (e.g., sufentanil; Vendruscolo et al. 2018) than for others. Interestingly, rats that were given the opportunity to self-administer sufentanil for 12 hours per day also shown somatic signs of opioid withdrawal (Vendruscolo et al. 2018).

Non-contingent chronic nicotine exposure and nicotine self-administration procedures have evolved over the years and more rapidly in recent years. For many years, chronic non-contingent nicotine exposure was achieved through osmotic mini-pumps (Jonkman et al. 2008; Torres et al. 2013) and nicotine self-administration was achieved using intravenous (i.v.) drug delivery protocols (Watkins et al. 1999; O’Dell & Koob 2007). Several years ago, a nebulization protocol was developed for “bubbling” of pure liquid nicotine and delivery of nicotine in the breathing air of rodents (Cohen & George, 2013), usually for very chronic periods (e.g., 12 hours per day over many days). Using that technique, chronic nicotine vapor inhalation led to escalation of i.v. nicotine self-administration in male Wistar rats (Gilpin et al. 2014). More recent work used ENDS to show that nicotine vapor inhalation increases spontaneous locomotion and reduces body temperature in male Sprague-Dawley rats, and that these effects are blocked by systemic injection of mecamylamine (Javadi-Paydar et al. 2019).

In this study, rats exhibited time-dependent somatic withdrawal signs that were higher following 1) exposure to higher concentrations of nicotine vapor and 2) precipitation of withdrawal with systemic mecamylamine injection. One exception is the absence of any difference in spontaneous withdrawal signs between the 20 and 40 mg/mL groups at 24 hours post nicotine exposure both at day 1 and day 10 (Fig. 4); however there is a large difference between these 2 groups in spontaneous withdrawal signs at 1, 2, 4 and 6 hours post nicotine exposure on day 10 (not tested on day 1; see Fig. 5). Interestingly, mecamylamine-precipitated withdrawal signs followed a U-shape in the 20 mg/mL group. It is possible that mecamylamine-precipitated withdrawal signs in this group peaked just after mecamylamine injection and faded with time, then increased at late time points when spontaneous withdrawal signs are maximal (Figures 5 & 6). Although somatic nicotine withdrawal symptoms in rats are universally evaluated with a behavioral observation scale (Malin et al. 1992), many past studies did not evaluate nicotine withdrawal over time, making it difficult to directly compare those results with ours. In many studies, nicotine withdrawal was assessed for a longer duration (30 min), at a single time point immediately after termination of nicotine exposure or days-to-weeks later, and in some cases with mecamylamine injection before the last prolonged (12-24 hours) nicotine exposure (see Matta et al. 2007 for a review). To the best of our knowledge, only a few studies have evaluated nicotine withdrawal using the brief (10-15 min) observational test initially proposed in Malin et al. (1992), and at similar post-nicotine time points to those used in our study. The spontaneous withdrawal scores exhibited over time by rats in our 80 mg/mL group following 10 days of nicotine vapor exposure were similar to the withdrawal scores observed after 1 week of osmotic mini-pump nicotine exposure in male Sprague-Dawley (3 mg/kg/day nicotine tartrate; Malin et al. 1992) and male Wistar rats (9 mg/kg/day nicotine tartrate; Epping-Jordan et al. 1998). More recent work used the “nebulization” protocol to evaluate spontaneous withdrawal symptoms in male Wistar rats 30 min & 16 hours after termination of 1 week of 8 hour/day nicotine exposure (flow rate = 15 LPM; George et al. 2010), and in male Wistar rats 8 hours after termination of 2 weeks of 12 hour/day nicotine exposure (flow rate = 5 LPM; Gilpin et al. 2014); withdrawal scores in those studies were similar to the spontaneous withdrawal scores we observed in the 20 mg/mL group at 1, 6 and 24 hours following the 10^th^ day of 1-hour nicotine vapor exposure.

In our study, blood-nicotine and blood-cotinine levels emerge and vary over time in a concentration-dependent manner following 1 or 10 sessions of non-contingent nicotine vapor inhalation. One discrepancy is the blood nicotine levels of the 80 mg/mL group that were lower than the 40 mg/mL group on day 1 at 30-60 min post nicotine exposure, and the blood nicotine levels of the 40 and 80 mg/mL groups on day 10 that overlapped in all the time point except at 15 min post-nicotine exposure. These discrepancies could be due to variable nicotine metabolism among rats (on average t1/2=45 min in rats; Matta et al. 2007), a ceiling on intrapulmonary nicotine uptake after a threshold concentration, or both. To the best of our knowledge there is no evidence of a ceiling on intrapulmonary nicotine uptake in rats (or other species) after a threshold concentration. In other cited papers, we did not find discrepancies between nicotine doses and nicotine blood levels measured; however in most studies nicotine was administered at a single dose - and not through ENDS- and blood nicotine levels were measured at a single time point, making it difficult to compare those results with our study. In general, the pattern of results observed here show that blood-nicotine levels peak early after termination of vapor exposure then gradually decrease, blood levels of the nicotine metabolite cotinine rise as blood-nicotine levels fall, and blood levels of cotinine are higher after repeated sessions than after a single session for the 80 mg/mL concentration. We believe the higher start point for blood cotinine levels in 80 mg/mL rats on day 10 relative to day 1 is due to the development of metabolic tolerance for nicotine. Because nicotine is absorbed faster after inhalation than after daily injections or osmotic mini-pump delivery (Matta et al. 2013), we speculate that metabolic tolerance for nicotine may be more rapid with ENDS than with other routes of administration. Relative to past work examining nicotine pharmacokinetics using other routes of systemic nicotine administration in rats, blood-nicotine levels we measured in the 80 mg/mL group during the 2 hours following the 1^st^ day of nicotine exposure were similar to those previously reported in male Wistar rats following an i.v. nicotine injection (0.2 mg free base nicotine/kg body weight), but lower than those reported following a s.c. nicotine challenge (1.0 mg free base nicotine/kg body weight) (Craig et al. 2014). Blood-nicotine levels after a s.c. nicotine injection (0.8 mg/Kg nicotine bitartrate) in male Sprague-Dawley rats were similar to those observed here in the 40 mg/mL group immediately after the 1^st^ day of nicotine exposure (Javadi-Paydar et al. 2019). Subcutaneous osmotic mini-pumps commonly deliver to rats 1—4 mg/kg/day nicotine tartrate, with 1.0 mg/kg/day resulting in stable plasma nicotine levels of approximately 25 ng/ml (see Matta et al. 2007 for a review). These blood-nicotine levels are similar to those we observed in the 20 mg/mL group, but lower than those we observed in the 40 and 80 mg/mL groups. In the “nebulization” study cited above (George et al. 2010), blood-nicotine levels in male Wistar rats at 0 and 2 hours after the 1^st^ day of nicotine exposure were similar to the blood-nicotine levels we assessed in the 80 mg/mL group during the 2 hours following the 1^st^ day of e-vape nicotine exposure. In another study that used an ENDS to deliver aerosolized nicotine to male and female Sprague–Dawley rats (Werley et al. 2016), blood-nicotine levels (at an unspecified time point following termination of nicotine vapor exposure) after 28 or 90 days of nicotine exposure (1 mg/L for 16 min/day) were similar to the blood-nicotine levels in our 40 and 80 mg/mL groups after 10 days of nicotine e-cigarette vapor. Interestingly, the chronic 16 min/day nicotine vapor exposure produced blood-cotinine levels similar to our 20 mg/mL group, whereas chronic 48 min/day nicotine vapor exposure produced blood-cotinine levels similar to the 40 and 80 mg/mL groups in our study. The concentration of nicotine in the vape solution used in Werley et al. (2016) work (1 mg/L) was much lower than our concentrations (20, 40 and 80 mg/mL). However, in Werley et al. (2016), rats were exposed to nicotine aerosol continuously for 16 or 48 minutes, while we delivered 30 discrete 3-sec puffs over a period of 60 minutes. Moreover, the Werley et al (2016) study described their inhalation exposure as “nose-only”, because animals were gradually habituated to nose-only restraint and were exposed to nicotine “puffs” at a very close distance, whereas our work used delivery of nicotine vapor to the entire cage.

In our study, only male subjects were employed. It was previously reported that female rats self-administer (i.v.) nicotine at higher rates than males, and sex differences have been reported for the anxiogenic, anorexic, antinociceptive and sedative effects of nicotine administered through mini-pumps or injections (see Matta et al. 2007 and Cohen and George 2013 for reviews), but recent studies using ENDS did not report sex differences (Werley et al. 2016; Smith et al. 2019 bioRxiv: 10.1101/797084; doi: https://doi.org/10.1101/797084). No sex differences were found in the blood nicotine and cotinine levels, body weight changes, food consumption and lung inflammatory responses following chronic nicotine vapor exposure through ENDS in adult Sprague-Dawley rats (Werley et al. 2016; 28 or 90 days of nicotine exposure [1mg/L] for 16/48/160 min/day), and a more recent study also did not report sex differences in behavioral and physiological outcomes measured in adult Wistar rats exposed to ENDS (Smith et al. 2019, bioRxiv: 10.1101/797084; doi: https://doi.org/10.1101/797084). Interestingly, it has been suggested that nicotine delivery via ENDS may have different effects on the brain and periphery (i.e., cardiovascular, respiratory, immune, metabolic systems) than cigarette smoking and other routes of nicotine administration (Milano et al. 2019).

Similar to other studies (Nguyen et al. 2016a, 2016b, 2017, 2018; Javadi-Paydar et al. 2018, 2019; Vendruscolo et al. 2018) that delivered drugs through ENDS to rodents, we did not measure the drug concentration in the air of the chambers and in the fur of rats, or conduct an epidermal tissue biopsy. While nicotine vapor exposure in a chamber does not exclude uptake through skin/fur absorption or grooming, such effects seems to be negligible compared to inhalation effects observed in rodents following nicotine whole-body vapor exposure through ENDS (see Milano et al. 2019 for a review). Presumably, it would take much longer for “dilute” nicotine to be absorbed through the fur/skin (see Turner et al. 2011 for a review concerning the routes of drug administration in laboratory animals) and we would expect slow nicotine absorption and a stable increase of blood nicotine levels over time. Even if it is not possible dissociate nicotine uptake through the fur/skin from grooming, if most nicotine was absorbed through grooming we would not expect a gradual decrease in the blood-nicotine levels over time as we have observed, since it is highly unlikely that all animals engaged in grooming behavior in such a temporally (and decreasing) organized manner. The fact that overall blood nicotine levels decreased over time seems to rule out the possibility that the majority of nicotine occurred through fur/skin absorption or grooming. However, whole-cage vapor delivery may still allow the possibility for drug absorption into the fur that can lead to time-dependent spikes in blood-drug and/or blood-metabolite concentrations if animals engage in grooming behavior during and after vapor exposure (we cannot exclude this as a possible explanation for the 80 mg/mL group at 120 min time-point on day 1, and for the 40 mg/mL at 30 min time-point on day 10, where blood-nicotine levels have increased in comparison to the previous time point). Future studies in the field should directly address these points with analysis of air-nicotine level, lung and epidermal tissue biopsy and determination of nicotine content in the fur.

The data presented here indicate that using ENDS to deliver nicotine vapor to male Wistar rats produced: 1) blood-nicotine levels similar to those reported in human smokers (10 to 50 ng/mL; see Matta et al. 2007); 2) blood-nicotine levels similar to those previously observed in rats following other routes of systemic nicotine administration; 3) somatic withdrawal signs comparable to those observed after chronic nicotine treatment using other routes of administration. Our data also provide parameters that can be used as a reasonable starting point for future work that employs non-contingent nicotine vapor inhalation in rats using ENDS. However, there are many parameters that can and should be manipulated in future work to identify the non-contingent exposure conditions that most closely approximate the human condition and also the parameters that are most likely to support the acquisition of an operant task for nicotine e-cigarette vapor inhalation. These hardware and session parameters may include but not be limited to changes in puff duration, vehicle composition (proportions of PG:VG), inclusion of flavorants in the vehicle, inter-puff time intervals, session duration, number of sessions, and exhaust settings. In our study, PG:VG ratio was selected based on the fact that e-cigarette liquid formulations used by humans are often composed of 50:50 PG:VG (Peace at al. 2016). However the impact of PG:VG ratio on nicotine use is also not yet clear, and one recent study shows that PG/VG ratio does not impact ENDS use (Smith et al. 2019). Future work should also examine the impact of age, sex, strain, and species on nicotine vapor effects on brain and behavior. Finally, ENDS can be used to examine a wide array of behavioral endpoints that have been tested with other methods of chronic nicotine vapor delivery: for example, self-administration of nicotine using other methods (e.g., i.v.; Gilpin et al. 2014), nociception (Baiamonte et al. 2013), and the effects of poly-drug use (e.g., nicotine and alcohol; Leao et al. 2015).

## Acknowledgments

Funding for this award was provided by National Institute of Drug Abuse (NIDA) Award R44DA046300 (MC & NWG). This work was supported in part by Merit Review Award #I01 BX003451 (NWG) from the United States (U.S.) Department of Veterans Affairs, Biomedical Laboratory Research and Development Service.

